# SeRenDIP-CE: Sequence-based Interface Prediction for Conformational Epitopes

**DOI:** 10.1101/2020.11.19.390500

**Authors:** Qingzhen Hou, Bas Stringer, Katharina Waury, Henriette Capel, Reza Haydarlou, Sanne Abeln, Jaap Heringa, K. Anton Feenstra

## Abstract

**Motivation:** Antibodies play an important role in clinical research and biotechnology, with their specificity determined by the interaction with the antigen’s epitope region, as a special type of protein-protein interaction (PPI) interface. The ubiquitous availability of sequence data, allows us to predicting epitopes from sequence in order to focus time-consuming wet-lab experiments onto the most promising epitope regions. Here, we extend our previously developed sequence-based predictors for homodimer and heterodimer PPI interfaces to predict epitope residues that have the potential to bind an antibody.

**Results:** We collected and curated a high quality epitope dataset from the SAbDaB database. Our generic PPI heterodimer predictor obtained an AUC-ROC of 0.666 when evaluated on the epitope test set. We then trained a random forest model specifically on the epitope dataset, reaching AUC 0.694. Further training on the combined heterodimer and epitope datasets, improves our final predictor to AUC 0.703 on the epitope test set. This is better than the best state-of-the-art sequence-based epitope predictor BepiPred-2.0. On one solved antibody-antigen structure of the COVID19 virus spike RNA binding domain, our predictor reaches AUC 0.778. We added the SeRenDIP-CE Conformational Epitope predictors to our webserver, which is simple to use and only requires a single antigen sequence as input, which will help make the method immediately applicable in a wide range of biomedical and biomolecular research.

**Availability:** Webserver, source code and datasets are available at www.ibi.vu.nl/programs/serendipwww/

## 1 Introduction

Protein-protein interactions (PPI) are crucial for most biological functions, and thus of great importance to understand cellular processes (Jones and Thornton, 1996). Therefore, an interest exists for discerning the mechanisms of PPI and discovering theoretical and practical applications such as in biological, biophysical and biochemical studies (Valencia and Pazos, 2002; Shoemaker and Panchenko, 2007; Gallet *et al*., 2000). Here we are particularly interested in the interactions between antibodies and their antigens, which can be considered as a specific form of PPI that has been shown to be very different from general interactions between proteins (Esmaielbeiki *et al*., 2016). The process of antibody-antigen-binding has gathered significant attention due to the high specificity and affinity of antibodies to their target (Sela-Culang *et al*., 2013). This property can be exploited in many areas, e.g. for the development of diagnostic tools, therapeutics, and peptide-based vaccines (Khan, 2014; Parvizpour *et al*., 2020), making antibodies one of the most important biopharmaceuticals today with an ever increasing amount of antibody-based therapies being approved for clinical use (Kaplon *et al*., 2020).

To facilitate the use of antibodies, it is vital to identify the distinct region on the antigen that will be recognised by an antibody; the specific amino acids of such a region are known as the *epitope* (Potocnakova *et al*., 2016). Identification of an antigen’s epitope regions and a better understanding of the mechanism of antibody-antigen-recognition will lead to improved antibody engineering and thus widen their applications in the future (Sela-Culang *et al*., 2013). Generally, there is a distinction between *continuous* (linear) epitopes and *discontinuous* (conformational) epitopes: a continuous epitope is comprised of a single continuous stretch of amino acids, while the residues forming a discontinuous epitope are made up of several stretches in the sequence that are brought together by the protein being folded (Barlow *et al*., 1986). Purely linear epitopes do not occur naturally but rather should be seen as segments of (larger) conformational epitopes (Rubinstein *et al*., 2008; Kringelum *et al*., 2013).

Large scale experimental epitope identification is not feasible as this process is time-consuming and costly (El-Manzalawy and Honavar, 2010). As a consequence, many efforts have been devoted to developing computational methods for prediction of epitopes instead, aiming to predict continuous or discontinuous epitopes, or both. Linear epitope predictors can be of use if the objective is to substitute a protein antigen by a peptide fragment in order to develop or produce antibodies, however, the desired cross-reactivity with the native protein antigen is often limited as almost all naturally occurring epitopes are conformational (Ponomarenko and Van Regenmortel, 2009). Continuous epitope prediction can also be applied to antibody detection based on denatured proteins deprived of their 3D structure (Sanchez-Trincado *et al*., 2017) and for the design of epitope-based vaccines (Parvizpour *et al*., 2020).

Discontinuous epitope prediction is more suitable to discover existing epitopes, i.e. to predict the explicit residues on the protein structure in its native fold that interact with an antibody. Such predictions can thus support immunodiagnostic and therapeutic methods that demand recognition of the natively folded protein (Forsström *et al*., 2015). Brown *et al*. (2011) showed that antibodies raised against full-length protein antigens consistently outperformed those raised against peptide antigens. Furthermore, prediction of conformational epitopes can help determine a list of epitope candidates to be confirmed by experimental testing (Sanchez-Trincado *et al*., 2017). An improved understanding of antibody-antigen-interactions also furthers our understanding of the immune response process in general (Zhang *et al*., 2011).

Epitope prediction tools can also be divided according to the input being structure- or sequence-based. Structure-based methods are usually able to outperform sequence-base methods in accuracy in direct comparison but are severely limited by the number of 3D protein structures available to use (Gao and Kurgan, 2014). Sequence-based methods offer the critical advantage of a large volume of data being available for training. Although the amount of protein structures accessible is also growing, there is still a lack of confirmed structural data for most antibody-antigen-complexes while the number of entries for protein sequences is increasing exponentially (Schwede, 2013). To take advantage of this vast amount of information, various different implementations of sequence-based epitope predictors have been developed over the last years. So far these methods have focused almost exclusively on linear epitope prediction (Saha and Raghava, 2006; El-Manzalawy *et al*., 2008; Davydov and Tonevitsky, 2009; Rubinstein *et al*., 2009; Sweredoski and Baldi, 2009; Ansari and Raghava, 2010; Wee *et al*., 2010; Yao *et al*., 2012; Gao *et al*., 2012; Singh *et al*., 2013; Shen *et al*., 2015; Jespersen *et al*., 2017; Liu *et al*., 2020), despite the uncommonness of naturally occurring linear epitopes.

In addition, we can also distinguish between fixed length and residue-specific epitope prediction methods. The first type, which is the majority of methods, outputs epitopes of a pre-defined length, typically between 12 and 22 amino acids. The second type of methods offers residue-specific prediction by assigning a score to each amino acid which quantifies its likelihood to be part of an epitope. Only a small number of methods provide residue specific predictions. AAPPred uses a support vector machine classifier based on amino acid pair frequency and antigenicity scales (Chen *et al*., 2007; Davydov and Tonevitsky, 2009). The model was trained on the linear epitope database Bcipep and randomly chosen non-epitopes taken from Swiss-Prot as well as experimental data. BepiPred-2.0 is currently the most widely used and cited method for epitope prediction (Jespersen *et al*., 2017). It was trained on antibody-antigen crystal structures taken from the Protein Data Bank (PDB) and applies a random forest algorithm to assign a probability score to every residue. Discontinuous epitope prediction from sequence is possible using BEST and CBTOPE (Ansari and Raghava, 2010; Gao *et al*., 2012; El-Manzalawy *et al*., 2017). As the field of sequence-based predictors of conformational epitopes is so limited, a strong need exists for new tools able to support the identification of the specific residues of a protein antigen interacting with its antibody. Here we focus on a predictor that can achieve accurate identification of residues that constitute the antigen’s epitope(s).

Previously, we developed a generic protein-protein interface predictor, that makes predictions on a per residue basis. This resulted in a widely usable interface predictor that only requires a single sequence as input (Hou *et al*., 2017, 2019). It is based on a random forest model and incorporates several features that can be derived from the sequence, including conservation (Pirovano *et al*., 2006; Hou *et al*., 2015), predicted secondary structure (Guharoy and Chakrabarti, 2007; de Vries and Bonvin, 2008) and solvent accessible area (Ofran and Rost, 2007; Li *et al*., 2012), and as novel features protein length (Hou *et al*., 2015, 2017) and predicted backbone flexibility (Cilia *et al*., 2013; Hou *et al*., 2017, 2019). Here we investigate if this approach using random forest models with previously selected features, and the combination of epitope data and general PPI data sets can help to make epitope prediction more accurate.

Antibody-antigen-interaction differs strongly from other PPI, hence if we want to use PPI prediction methods for the purpose of epitope prediction adaptation is required (Yao *et al*., 2013; Esmaielbeiki *et al*., 2016). In order to determine which type of data should be included to re-train our prediction approach, we first investigated the predictive performance of RF models previously trained on homodimer and heterodimer PPI datasets (Hou *et al*., 2015, 2017). We then assembled a new antibody-antigen-interface dataset, specifically using conformational epitopes derived from structural data to optimize the prediction of this type of epitope by the model. This data set was used to train our new epitope predictor SeRenDIP-CE: SEquence-based RanDom forest Interface Predictor for Conformational Epitopes, which aims to predict residues that have the potential to bind an antibody, and therefore be part of an epitope.

## 2 Materials and Methods

For clarity, we here summarize the training and testing protocols used to derive our random forest classifiers, following the procedure developed previously (Hou *et al*., 2017, 2019), and also the details of constructing the Dset_anti dataset for sequence-based epitope prediction.

### 2.1 Dataset

The SAbDab structural antibody database from the Oxford Protein Informatics Group (OPIG) (Dunbar *et al*., 2014) was used to obtain the antibody-antigen structures as our starting dataset. 2 023 PDB (Protein Data Bank) structures of antibody-antigen complexes were selected. The sequence of each antigen chain was extracted from PDB files. CDhit (Li and Godzik, 2006) was used to remove redundancy among all antigen sequences using 25% sequence identity (seq. ID) cut-off to obtain a non-redundant dataset of 311 antigen sequences. To evaluate the performance of our homodimer and heterodimer predictors developed previously, we further removed redundant sequences at 25% seq. ID between the antigen dataset and our homodimer and heterodimer datasets, retaining 280 antigen sequences (75 845 residues) as our antigen dataset (Dset_anti). To further validate the ability of our approach on epitope prediction, the validation set of 5 conformational epitopes from BepiPred-2.0 (Jespersen *et al*., 2017) was used as an independent test set.

For a fair comparison with other state-of-the-art predictors, we download the training set of conformational epitopes from BepiPred-2.0 (Jespersen *et al*., 2017) and removed sequences from our five test datasets redundant with their training set at a cutoff of 25% seq. ID. The number of proteins retained in each test set are reported in SI Table S1.

### 2.2 Generation of protein sequence features

For each antigen sequence, we obtain a series of features per position across the query sequence, as previously described (Hou *et al*., 2017, 2019). In short, for each input sequence, PSI-BLAST (version 2.2.22+; Altschul *et al*., 1997, 2005; Schäffer *et al*., 2001) is used to retrieve sequence homologs from the NR70 database using max. 3 iterations, an E-value threshold of 10^*−*5^ and max. 500 hits. Multiple Sequence Alignments (MSAs) of the query sequence and its PSI-BLAST hits were aligned using Muscle (Edgar, 2004), and profiles for each of the hit sequences were generated by re-mastering from the Multiple Sequence Alignments (Hou *et al*., 2019). The resulting profiles were then used as input to NetSurfP to predict solvent accessibility and secondary structure. Sequence entropy values are calculated at each column of the alignments to quantify the conservation for each position of the query sequence. Backbone flexibility scores was predicted using Dynamine (Cilia *et al*., 2013).

For each type of feature, in addition to the feature value for the query position, the average and standard deviation for the homologs in the alignment are also used, resulting in 20 features. A fixed nine residue sliding window is implemented to include the same features from neighbouring residues. All features used are listed in SI Table S3, for more detail, please refer to Hou *et al*. (2017).

### 2.3 Definition of epitope residues

Epitope residues were defined based on the distance between atoms in the antibody and antigen; when this is less than 6.0 Å it is assumed that the antigen residues are interacting with the antibody. This approach is used for all 280 antigens, resulting in 7 147 epitope residues and 68 698 non-epitope residues.

### 2.4 Training procedure

To obtain a reliable and stable prediction, we randomly split 280 proteins of the antigen dataset Dset_anti into 80% training (224 sequences) and 20% testing (56 proteins) datasets and repeated this five times, retaining five coupled training and test datasets. Five predictors were generated using each of the training sets separately. Note that we repeat the splitting process to avoid over-fitting and biases in the training datasets and to ensure stable training of our predictors. The cross-validation was done on each training dataset during the training process.

Random forest is used to integrate all epitope related features derived and predicted from the antigen sequences to generate the epitope predictor. The Random Forest R-package (Liaw and Wiener, 2002) was used to construct predictors and test performance. The number of variables randomly sampled at each split of the forest is defined by the global parameter ‘mtry’; the value with the best AUC-ROC (see below under Benchmarking) was selected by a grid search over a set interval of values from one to 20. 10-fold cross validation of the training dataset was implemented, with one fold used to find the best ‘mtry’. The cross-validation and random forest parameter tuning were done using the caret R-package (Kuhn, 2015). The final model was then fitted on the whole training dataset using the fixed best ‘mtry’. The 10-fold cross validation was done 3 times with random seeds. The ratio of epitopes to other positions is about 1:10 (7 147 vs 68 698), which we balanced by downsampling the majority class (non-epitope) in our training dataset to ensure the same frequency (1:1) between the classes. Downsampling is suggested by others (Lin and Chen, 2013) and also outperformed oversampling in our previous work (Hou *et al*., 2017).

Feature importance was measured by using the MeanDecreaseGini function in the R ‘caret’ package, which measures variable importance based on the Gini impurity index (Kuhn, 2015).

### 2.5 Testing procedure

After obtaining the predictors, the testing procedure was performed by applying the predictor on its corresponding test set. The average and standard deviation of the performances metrics (see below under ‘Benchmarking’) were computed for comparison with other approaches using the predictors trained on the five datasets. We also evaluated the performance of our previously derived homodimeric and heterdimeric predictors on our epitope Dset_anti test sets.

To further validate the ability of our approach on epitope prediction, the conformational epitope evaluation set of five antigen proteins from BepiPred-2.0 (Jespersen *et al*., 2017) was used as another independent test set. For a fair comparison, for each protein only predictions were made using models with no redundant sequences in their training set at 25% Seq. ID.

### 2.6 Benchmarking

We evaluate the performance of our trained models on the test sets, using the usual measures as used before (Hou *et al*., 2017, 2019). Here, true positives (TP) are correct prediction of epitope residues, false positives (FP) are residues incorrectly predicted as epitope, true negatives (TN) designate non-interacting sites that were recognised, and false negatives (FN) are non-epitope sites which were predicted as epitope.

- Precision (Positive Predictive Value, PPV) = TP / (TP+FP)
- Recall (True Pos. Rate, TPR, Sensitivity, Coverage) = TP / (TP+FN)
- F1 = 2 *×* Precision *×* Recall / (Precision + Recall)
- Specificity (True Negative Rate) = TN / (TN+FP); Error (1*−*Specificity, False Positive Rate, FPR) = FP / (TN+FP)
- Accuracy = (TP+TN) / (TP+FN+TN+FP)
- AUC-ROC: area under the cuve of the ROC plot (see below)

We furthermore explored the models’ predictions in more detail using Receiver-Operator Characteristics (ROC; TPR vs. FPR) and Precision/Recall (P/R; precision vs. recall) plots.

## 3 Results

To benchmark epitope prediction, we extracted antigen sequence and structure data from SabDab, yielding 280 antigens whose epitopes are determined using PDB structures. These we divided into training and test sets randomly for five times, resulting in five pairs of training and test sets which comprise our new Dset_anti. We followed the procedure developed previously (Hou *et al*., 2017, 2019), as summarize above in Methods. In total, 172 features derived from sequences are used including conservation, solvent accessibility, flexibility and secondary structure. We then compare prediction performance among random forest models trained on these features for three types of interface data: homodimers, heterodimers and epitopes.

### 3.1 Generic PPI prediction to detect Epitope interface

We first evaluated the performance of our existing generic PPI predictors to investigate how well these may function for epitope prediction. Table 1 shows the prediction performance of different predictors trained on generic homodimer, heterodimer and combined PPI datasets. On the Dset_anti test sets, the heterodimer predictor obtained the highest AUC-ROC (0.666) compared to the other two. The combined homo/hetero predictor, which performed well on the heteromeric test set (AUC-ROC 0.655, Hou *et al*., 2019), does not provide good predictions for the epitope test sets (AUC 0.585). Also the homodimer predictor preforms quite poorly (AUC 0.555). From this we could infer that heterodimer and epitope interfaces share some common patterns, while in contrast, adding homodimer data to the training seems to bring noise into the epitope prediction.

**Table 1.** Performance of our previously developed generic interface predictors on epitope prediction, as measured by accuracy, sensitivity, precision, specificity, F1 and AUC-ROC vs. the Dset_anti test sets in five-fold validation; in all cases higher is better. Highest values per metric and those less than one standard deviation below the highest are indicated in bold.

| Training set(s) |  | Acc | Sens | Prec | Spec | F1 | ROC |
| --- | --- | --- | --- | --- | --- | --- | --- |
| Hetero | Average | **0.750** | 0.439 | **0.170** | **0.781** | **0.245** | **0.666** |
|  | St.Dev | 0.035 | 0.024 | 0.007 | 0.038 | 0.008 | 0.026 |
| Homo | Average | 0.529 | **0.550** | 0.106 | 0.526 | 0.177 | 0.555 |
|  | St.Dev | 0.021 | 0.017 | 0.011 | 0.023 | 0.015 | 0.017 |
| Homo+Hetero | Average | 0.558 | **0.558** | 0.114 | 0.558 | 0.189 | 0.583 |
|  | St.Dev | 0.026 | 0.013 | 0.010 | 0.028 | 0.014 | 0.022 |

### 3.2 Epitope prediction using different training sets

After checking the performance of the previously trained generic interface predictors (Hou *et al*., 2017, 2019), we combined the antigen and homo- and heteromeric datasets to further develop a new epitope predictor. Table 2 show the epitope prediction performance by predictors trained on the antigen dataset and in combination with the homodimer and heterodimer datasets. The ‘Antigen’ predictor performs well on epitope prediction, yielding AUC-ROC of 0.694. Adding the homodimer dataset for training does not increase the prediction performance, similar to what was observed for the ‘generic homo’ PPI prediction (Table 1). Combined training including all three datasets also failed to improve epitope prediction. Interestingly, the best performance could be obtained from the models trained on antigen and heterodimer datasets whose AUC-ROC reaches 0.704, which is consistent with high performance for epitope prediction of the generic ‘hetero’ PPI predictor as shown in previous work (Hou *et al*., 2017, 2019).

**Table 2.**
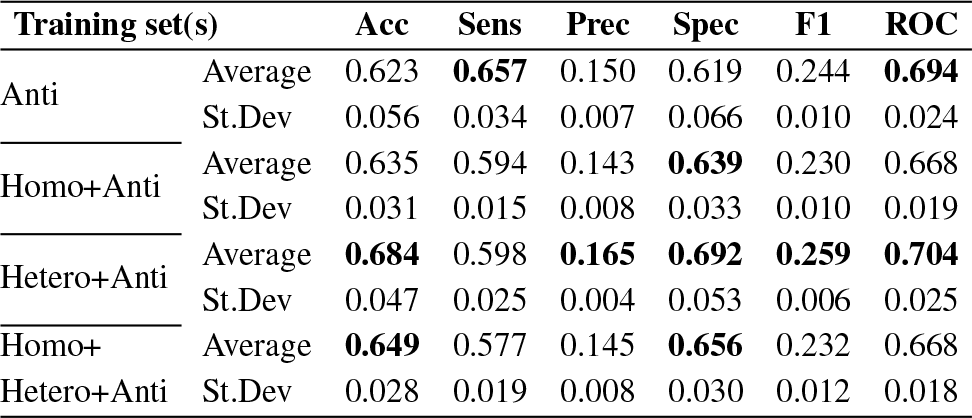
Performance of SeRenDIP-CE epitope predictors trained on the Dset_ anti training sets and in combination with our existing homo- and heteromeric datasets vs. the Dset_anti test sets. Measures and other details as in Table 1.

### 3.3 Feature importance

We considered which features were most informative for the final epitope prediction. As previously observed for our generic PPI predictors (Hou *et al*., 2017, 2019), sequence length, NetSurfP predicted accessibility of the query sequence and their mean values across homologs along the alignments are the leading contributions also for epitope prediction, as shown in SI Fig. S1 which shows the importance of all 172 features by ‘mean decrease GINI’ on the random forest models.

Note that, the Dynamine predicted flexibility scores were important to predict homodimer and heterodimer interface residues, but apparently less so for epitope prediction. Conversely, Entropy as a measure for conservation is most important for epitope prediction, less crucial for heteromeric interfaces, and least for homomeric interface prediction. Apparently, generic homodimeric interfaces differ from epitope interfaces in terms of flexibility and conservation.

To confirm the surprisingly high importance of length as a global feature for predicting local epitope residues, and to assess its relative importance compared to solvent accessibility, we also built models excluding length or/and solvent accessibility from the model features. The AUC-ROC drops from 0.70 to 0.66 without solvent accessibility. When just length is removed the performance drops even more to 0.63. The performance of the predictors that excluded both length and accessibility features dramatically drops to 0.59 which shows the importance of length and accessibility (see SI Table S2 for details). We also map the AUC of each protein onto the sequence length, showing only a weak correlation (*r* = 0.25) between the sequence length with the prediction performance (see SI Fig. S2); hence, our predictors show a similar predictive power for antigens independent of their length.

### 3.4 Comparison with other sequence-based epitope predictors

To show that our SeRenDIP-CE models capture relevant properties of epitope interfaces, we compared our predictors with other published and available state-of-the-art sequence-based epitope prediction methods: BepiPred-1.0 and -2.0 (Larsen *et al*., 2006; Jespersen *et al*., 2017), and AAPPred (Davydov and Tonevitsky, 2009). The comparison is done using our Dset_anti test sets filtered at 25% seq. ID w/r to the BepiPred training set. Our models trained on Anti and Anti+hetero obtain highest AUC-ROC (0.716 and 0.728 resp., see SI Table S1) compared to the other methods (around or below 0.6), and in the ROC plot (Figure 1) both achieve consistently higher coverage (TPR) for any error (FPR). Furthermore, the combined predictor (hetero+antigen) achieves overall high precision in the P/R plot (Figure 1), compared to the other approaches tested, while at a low recall, the ‘antigen’ predictor obtains highest precision.

**Fig. 1.**
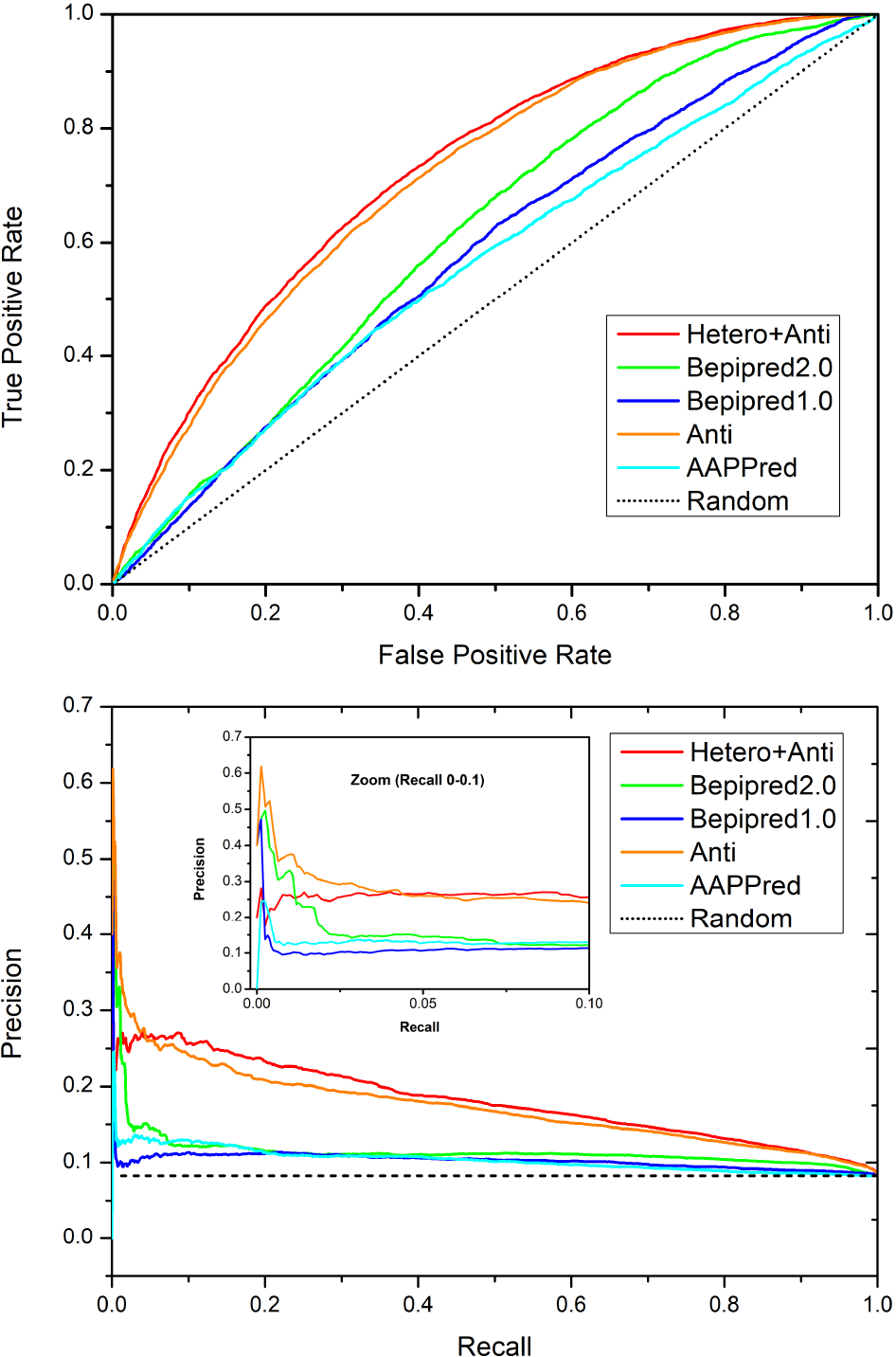
Comparison with other approaches by ROC (top) and P/R (bottom) plots. Red (combined ‘hetero/antigen’) and orange (‘antigen’) represent our two approaches; blue, green and cyan represent state-of-the-art BepiPred-1.0, 2.0, and AAPPred, respectively; and black represents random performance. In both analysis we can observe that, compared to the other methods, both SeRenDIP-CE models consistently obtain higher recall for a given error in the ROC plot, and higher precision for a given recall in the P/R plot; at low recall, shown in the inset for recall 0.0 *−* 0.1, BepiPred-2.0 and our specific ‘Antigen’ trained model both achieve highest precision.

We furthermore compared our predictors with BepiPred-2.0 on their independent conformational epitope dataset (Jespersen *et al*., 2017). As can be seen from Table 3, with an average AUC-ROC of 0.683 and 0.652, our two predictors perform better than BepiPred-2.0 at 0.603, which was previously reported to be the best conformational epitope predictors on their five-protein validation set (Jespersen *et al*., 2017).

**Table 3.**
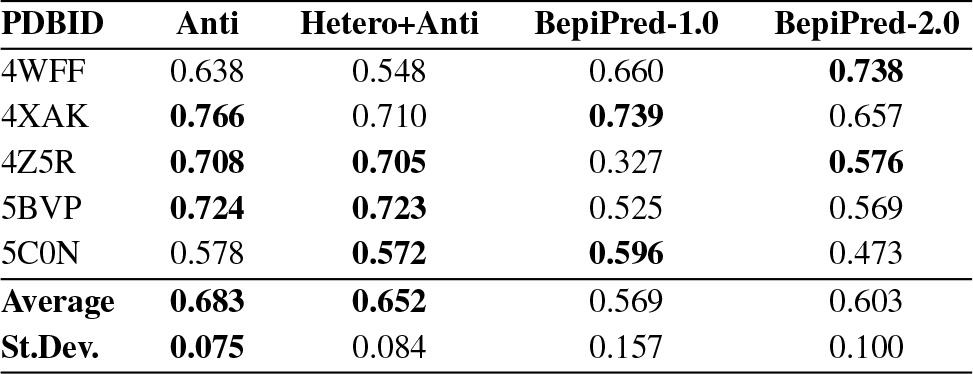
Performance in AUC-ROC of our epitope predictors on the independent conformational epitope dataset from BepiPred-2.0 (Jespersen et al., 2017), and comparison with BepiPred-1.0 and -2.0. Scores within one standard deviation from the maximum per protein and the overall average are indicated in bold. The lowest standard deviation is also indicated in bold.

### 3.5 Application to COVID-19 virus RNA Binding Domain

We further evaluate our approach to the COVID-19 virus spike receptor-binding domain against the solved crystal structure complexed with a neutralizing antibody (PDB:7BZ5). The spike receptor-binding domain (chain A) was used as input sequence by our approach to compute the probability score of each amino acid being an epitope. The structural details of its structure (in cyan) and epitope region (in red) can be seen in Figure 2.A. Our approach could capture most of the epitope positions solved in 7BZ5 at high probability (red in Figure 2.B and in C track ‘Prediction (%)’) and the majority of false positives locate around the real epitope (red in 2.C track ‘7BZ5’). Note that our predictor is not anti-body specific, hence one should interpret the predictions to show which residues have the *potential* to bind to an antibody. In this context, our approach is quite accurate with an AUC-ROC of 0.78 an a high precision of 56% at low recall (12%; see SI Fig. S4 for the corresponding ROC and P/R plots).

**Fig. 2.**
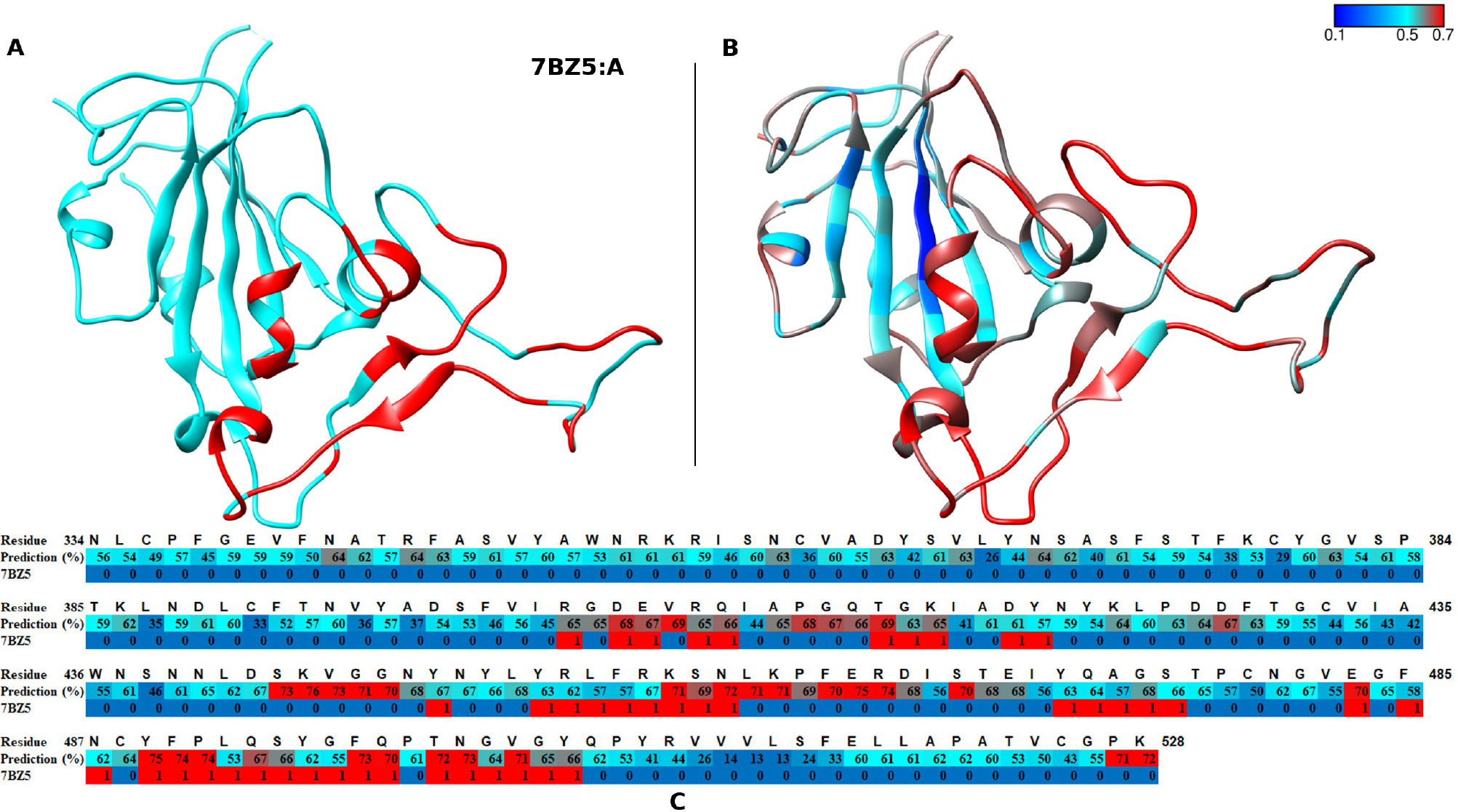
Epitope prediction of COVID-19 RBD protein. (A) The structure of COVID19 virus RBD (in cyan) and its epitopes (red). The full structure of antibody-antigen complex (7BZ5) is shown in SI Fig. S3. (B) The prediction performance of our predictor. The color gradient from blue to cyan to red shows the increased probability scores of that position being an epitope. (C) The details of the probability score at each position. Each block shows for a given residue from top to bottom: the residue number, the predicted score and the annotation if the residue is an epitope (1, red) or not (0, blue) according the PDB structure 7BZ5. The color scheme of the predicted score is as the same in the structure (B).

### 3.6 Showcase – Adiponectin receptors-2

To highlight the impact of accurate epitope prediction, we also investigated the Adiponectin receptor-2 bound to antibody fragments (PDB:5LX9). For this particular interaction, a coverage of 58% of 24 epitope sites at an accuracy of 82% is achieved, with an AUC-ROC of 0.819. Details can be seen in SI Fig. S5. Interestingly, our false positive predictions clustered on the other side of protein which might be due to the fact that the other side of transmembrane protein is also amenable to antibody binding.

### 3.7 The Webserver

We extended our SeRenDIP-CE webserver (www.ibi.vu.nl/programs/serendipwww/; Hou *et al*., 2017, 2019) to include the new epitope interface predictors presented in this work. The webserver is simple to use and is aimed at non-experts in both academia and industry. The input is only the antigen protein sequence. The predictions are based on the average of the five-fold trained models; for each position in the input sequence the average model score is reported, representing the predicted likelihood of being an epitope position, as well as the standard deviation to allow an estimate of the significance of the prediction.

## 4 Discussion

Epitope prediction based on antigen sequence is still a difficult problem, nevertheless our results show that the newly developed method SeRenDIP-CE is able to make significant progress in this field: the predictors reach an AUC-ROC of around 0.7. Note that, our epitope predictions effectively indicate which residues have the potential to bind an antibody, even though our data set consists of specific antigen-antibody complexes. Such a dataset does not necessarily cover all possible antibody binding sites, which will lead to an underestimation of the number of true positives.

Interestingly, the predictors trained on the combination of heterodimer and antigen datasets obtain the best performance. Indeed, the epitope interactions may be seen as a special type of heterodimer and therefore may be expected to share some common properties which could be used for the epitope prediction. To the best of our knowledge, this is the first time heterodimer data is used to improve the prediction of epitope positions. That might open a possibility of implementing ‘combined’ datasets to boost prediction accuracy of a particular ‘subset’ interface type of interest. Further improvement could come from applying deep learning algorithms or implementing additional properties as features, e.g. co-evolution signals derived from large sequence alignments, now that we have shown the power of careful selection of the datasets used for training.

Sequence based prediction is the only option when there is no structure or suitable template available, e.g. as may well be the case for the proteins in a newly discovered virus. Our method performs well on the epitope prediction of a COVID19 antibody-antigen complex. As can be seen in Figure 2, the false positives cluster around the epitopes annotated from the 7BZ5 structure. Given the high overall performance of our approach, predicted positions with high probability score might be interesting candidate targets for antibody design. Especially for virus proteins whose structures are not solved at the moment, one could expect to implement our method to improve selection of candidate epitope regions.

## 5 Conclusion

Our SeRenDIP-CE Sequence-based Random forest Epitope Predictors outperform other state-of-the-art approaches. This shows that co-training on epitope data and heteromeric PPI interface data, has the potential to make epitope prediction more robust. The webserver based on our predictor is simple to use and only needs one sequence of the protein of interest as input. We therefore expect that the SeRenDIP-CE method will be immediately applicable to a wide range of biomedical and biomolecular problems.

## Acknowledgements

KW and SA have received funding from the European Union’s Horizon 2020 research and innovation programme under the Marie Skłodowska-Curie grant agreement No 860197, the MIRIADE project. The authors declare that there is no conflict of interest.

