## Supplemental Material for "SeRenDIP-CE: Sequence-based Interface Prediction for Conformational Epitopes"

Bioinformatics
doi.10.1093/bioinformatics/xxxxxx

Supplementary Material

### SeRendEP: Sequence-based Interface Prediction for Conformational Epitopes

**Qingzhen Hou, Bas Stringer, Katharina Waury, Henriette Capel, Reza Haydarlou, Sanne Abeln, Jaap Heringa, K. Anton Feenstra**

www.ibi.vu.nl/programs/serendipwww/

1. **Comparison of different methods on antigen test sets**
2. **Prediction performances with and without sequence lengh and/or accessibility**
3. **List of proteins in the antigen dataset**
4. **Feature importance of the antigen and combined predictors**
5. **The prediction performance vs the sequence length**
6. **The structure of COVID19 RBD domain and it antibody solved in PDB 7BZ5**
7. **The prediction performance on COVID19 RBD domain in ROC plot and Precision-Recall plot**
8. **Epitope prediction on Adiponectin receptors-antibody complex**

**SI Table S1.** Comparison of different methods on antigen test sets. The AUC of ROCs of each method are shown in the table. The numbers of sequences in each datasets are also listed. Per method, AUC-ROC and average values that are within one standard deviation from the maximum are indicated in bold.

| Dataset | # seq | Aapred | Anti | Anti+ hetero | BepiPred 1.0 | BepiPred 2.0 |
|---|---|---|---|---|---|---|
| test1 | 31 | 0.515 | **0.693** | **0.715** | 0.550 | 0.590 |
| test2 | 37 | 0.591 | **0.721** | **0.732** | 0.575 | 0.628 |
| test3 | 33 | 0.567 | **0.755** | **0.766** | 0.562 | 0.630 |
| test4 | 33 | 0.541 | **0.684** | **0.696** | 0.585 | 0.593 |
| test5 | 34 | 0.597 | **0.725** | **0.733** | 0.589 | 0.641 |
| Average | 33.6 | 0.562 | **0.716** | **0.728** | 0.572 | 0.616 |
| Std.dev. | | 0.034 | 0.028 | 0.026 | 0.016 | 0.023 |

**SI Table S2.** Prediction performances with and without sequence length (Len) and/or accessibility (Acc) features predicted from sequence. The predictors were trained based on the antigen training sets. The performance was averaged by scores on five testing datasets as described in the main text. Per measure, highest scores are indicated in bold.

| Acc | Sens | Prec | Spec | NPV | F1 | Kappa | ROC | +Std.dev | Features |
|---|---|---|---|---|---|---|---|---|---|
| 0.623 | **0.657** | **0.150** | 0.619 | **0.947** | **0.244** | **0.110** | **0.694** | **0.024** | All |
| 0.546 | 0.646 | 0.124 | 0.536 | 0.937 | 0.208 | 0.063 | 0.634 | 0.018 | no Len |
| **0.648** | 0.586 | 0.147 | **0.653** | 0.939 | 0.235 | 0.103 | 0.675 | 0.022 | no Acc |
| 0.593 | 0.467 | 0.107 | 0.606 | 0.918 | 0.174 | 0.028 | 0.565 | 0.019 | no Len+Acc |

© The Author 2020.

**SI Table S3.** The list of 280 PDB IDs used in the Dset_anti antigen dataset.

| | | | | | |
|---|---|---|---|---|---|
| 1AHW_C | 2XWT_C | 4CKD_A | 4XWG_A | 5MJE_A | 6C5W_A |
| 1AR1_B | 2YBR_C | 4DKE_A | 4Y7M_C | 5N7W_X | 6C9U_A |
| 1BGX_T | 2ZJS_Y | 4EDW_V | 4YBQ_A | 5N88_E | 6CMG_A |
| 1BZQ_A | 3AB0_A | 4ETQ_C | 4YZF_A | 5NH3_A | 6CSE_C |
| 1EGJ_A | 3B9K_B | 4F2M_E | 4ZFG_A | 5NJ3_A | 6CW3_F |
| 1EZV_E | 3BT2_U | 4F37_A | 4ZSO_E | 5NMV_K | 6CYF_A |
| 1FE8_A | 3CFI_B | 4F3F_C | 5B3J_C | 5NUZ_C | 6DJP_A |
| 1FJ1_E | 3EZJ_A | 4G7V_S | 5BO1_A | 5O0W_A | 6DZM_D |
| 1FNS_A | 3FMG_A | 4GFT_A | 5C3L_C | 5O14_A | 6E1K_A |
| 1FSK_A | 3G6D_A | 4GRW_A | 5C6T_A | 5O1R_A | 6ELU_A |
| 1JRH_I | 3GI8_C | 4HCR_A | 5C8J_I | 5O2U_A | 6EQI_C |
| 1KB5_A | 3IU3_I | 4HT1_T | 5CBA_E | 5OB5_A | 6EY0_A |
| 1KXQ_A | 3JBA_A | 4HWB_A | 5CZV_A | 5OGI_A | 6EY6_A |
| 1LK3_A | 3JCX_A | 4I9W_A | 5D1Q_E | 5OVW_A | 6F2G_A |
| 1NFD_B | 3K81_C | 4IJ3_A | 5D1Z_I | 5SY8_O | 6F5G_A |
| 1NMB_N | 3KR3_D | 4IOF_A | 5D8J_A | 5TD8_C | 6FE4_A |
| 1NSN_S | 3KS0_A | 4JQI_A | 5DA0_A | 5TH9_A | 6FV0_A |
| 1OB1_C | 3LD8_A | 4JR9_A | 5DFV_A | 5TIH_A | 6FXN_A |
| 1OP9_B | 3LEV_A | 4JZJ_C | 5DFZ_C | 5TJW_A | 6GCI_A |
| 1QFW_A | 3LH2_S | 4KFZ_A | 5DMI_A | 5TQ0_A | 6GKU_A |
| 1QFW_B | 3LHP_S | 4KML_A | 5DWU_A | 5TZ2_C | 6GS1_A |
| 1SY6_A | 3LIZ_A | 4LMQ_D | 5E0Q_B | 5UKB_A | 6GV1_A |
| 1T03_B | 3MJ9_A | 4LQF_A | 5E7F_G | 5USF_A | 6GV4_B |
| 1V7M_V | 3MXW_A | 4LVN_A | 5ESZ_C | 5UTZ_A | 6H02_A |
| 1WEJ_F | 3NH7_A | 4M3K_A | 5F72_C | 5VKD_A | 6H16_A |
| 1XCQ_P | 3O0R_B | 4NP4_A | 5F93_A | 5VNW_A | 6H1F_B |
| 1YJD_C | 3OGO_A | 4NZR_M | 5FB8_C | 5VOB_D | 6H3T_A |
| 1YNT_F | 3OPZ_A | 4O9H_A | 5FHX_A | 5VOD_C | 6I04_A |
| 1Z3G_A | 3PGF_A | 4OII_A | 5FOJ_B | 5VXK_A | 6I07_C |
| 2A0L_A | 3PJS_K | 4OKV_E | 5H30_A | 5VYF_C | 6I6J_A |
| 2ARJ_Q | 3PNW_C | 4PLJ_A | 5H35_C | 5W5X_A | 6I9I_C |
| 2B2X_A | 3R1G_B | 4QEX_A | 5H8O_B | 5WTH_B | 6IBL_A |
| 2EXW_A | 3RAJ_A | 4QO1_B | 5HDQ_A | 5X0T_E | 6IDI_D |
| 2I9L_I | 3RU8_X | 4RDQ_A | 5I6X_A | 5XBM_C | 6MB3_E |
| 2J88_A | 3SKJ_E | 4RGM_A | 5IKC_M | 5YHL_A | 6MEI_C |
| 2JEL_P | 3TT1_A | 4RRP_M | 5IP4_D | 5ZS0_C | 6MUI_B |
| 2LTQ_A | 3U2S_C | 4U0R_A | 5J13_A | 6A0Z_A | 6N51_A |
| 2QQK_A | 3U30_A | 4UAO_A | 5J1S_A | 6A3W_C | 6NB3_A |
| 2R56_A | 3U9P_C | 4W2O_B | 5JMO_A | 6AJ7_C | 6NFJ_A |
| 2UZI_R | 3UBX_A | 4W6W_A | 5JQ6_A | 6AL5_A | 6NJL_A |
| 2VXQ_A | 3UX9_A | 4WEU_A | 5K59_A | 6APD_A | 6NN3_A |
| 2VXT_I | 3V7A_A | 4WUU_A | 5KEL_A | 6AVQ_B | 6O3B_C |
| 2VYR_A | 3W9E_C | 4XI5_B | 5KOV_A | 6B0N_G | 6OTC_A |
| 2WZP_A | 3X3F_A | 4XMM_D | 5KTE_A | 6BDZ_A | 6QEE_A |
| 2X7L_M | 4AG4_A | 4XMN_E | 5LX9_A | 6BGT_C | 6R7T_B |
| 2XQB_A | 4CAD_C | 4XT1_A | 5M30_A | 6BPA_A | 2XRA_A |
| 4CDG_A | 4XTR_A | 5MHR_A | 6C5V_A | | |

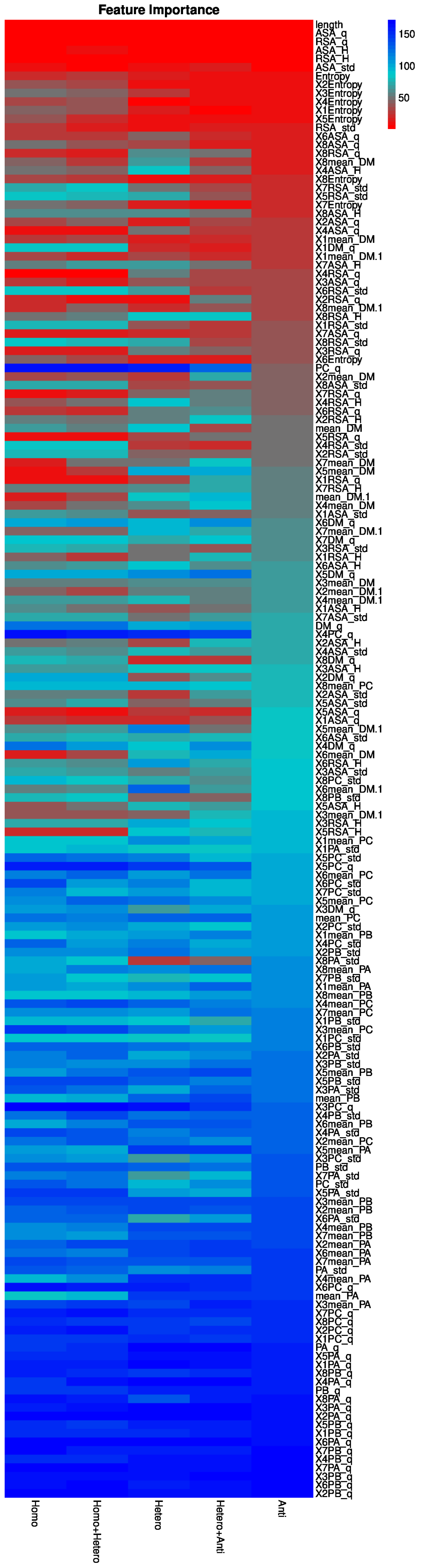


**SI Figure S1.** Heatmap of feature importance of generic homodimeric, heterodimeric, homo+hetero, Antigen and Anti+hetero predictors. More red, the more importance of that feature. The bluer, the less important the feature is. The features is ranked by the average importance of predictors trained by antigen dataset.

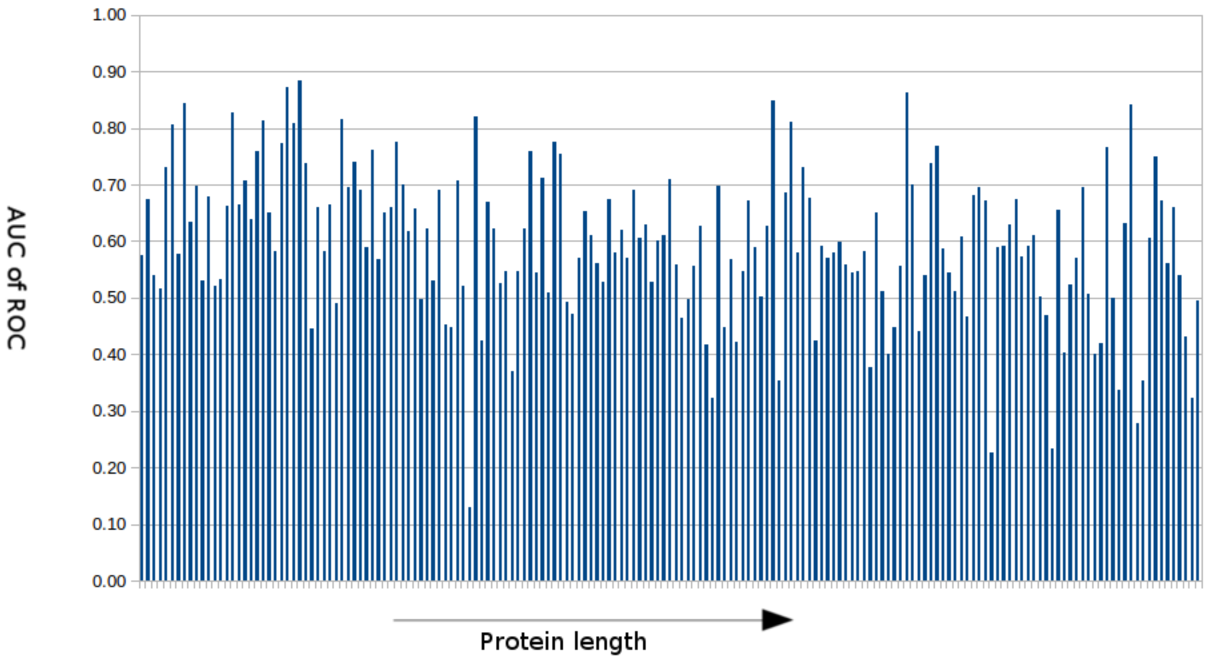


**SI Figure S2.** The prediction performance of our predictor vs. the sequence length in the testing dataset. The X axis shows the protein length. The length decreases from the left to the right.

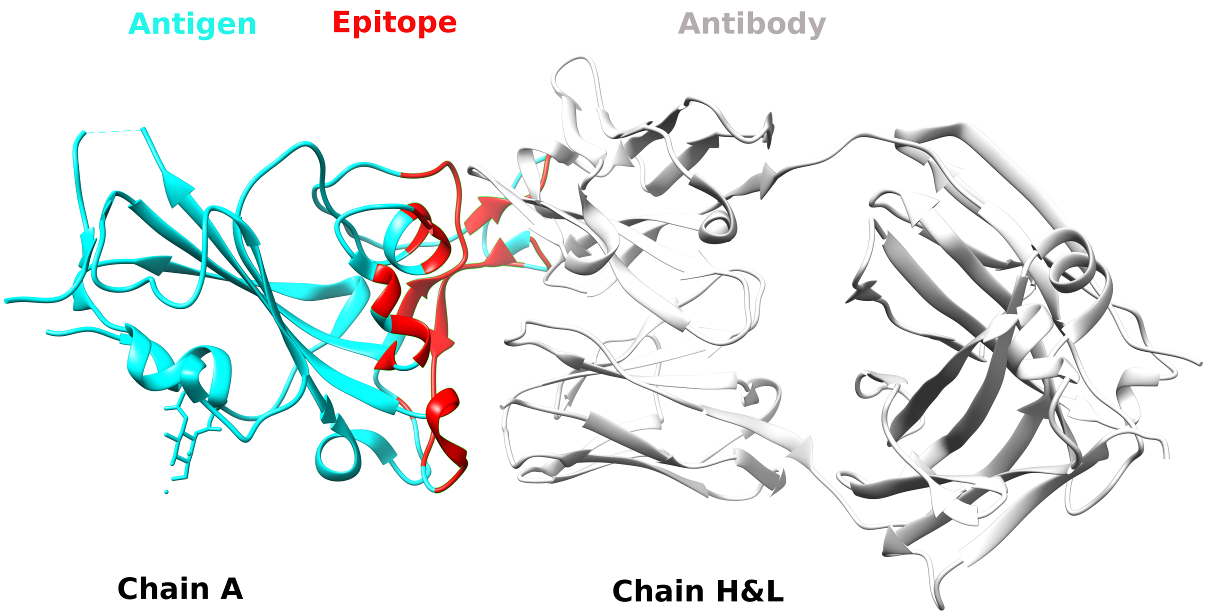


**SI Figure S3.** The structure of COVID19 RBD domain and it antibody solved in PDB 7BZ5. The antigen (RBD, 7BZ5:A) was shown in cyan and the epitopes are in red. The heavy and light chain (7BZ5:H&L) of the antibody are in white.

**SI Table S4.** In total, 6 feature types were used: Dynamine score, RSA, ASA, secondary structure, Entropy, and the length of query sequence. Average and standard deviations are calculated across the alignment of PSI-BLAST hits from the query sequence. For local features (Dynamine score, RSA, ASA, secondary structure and Entropy), 9-residue windowing approaches are used, which are indicated by a pre-pended 'X' and number (e.g. X1ASA_H), where $1 \cdots 4$ correspond to position $i-4 \cdots i-1$ and $5 \cdots 8$ to $i+1 \cdots i+4$.

| Name | Description |
|---|---|
| **DynaMine** | **Predicted backbone dynamics** |
| DM_q | the DynaMine score for the query sequence |
| mean_DM, sd_DM | the average and standard deviation of DynaMine score for each column in the alignment |
| **RSA/ASA** | **NetSurfP predicted Relative and Absolute Surface Accessibility (RSA, ASA)** |
| RSA_q | predicted RSA for the query sequence |
| RSA_H, RSA_std_H | average and standard deviation of predicted RSA for each column in the alignment |
| ASA_q | predicted ASA for the query sequence |
| ASA_H, ASA_std_H | average and standard deviation of predicted ASA for each column in the alignment |
| **Sec. Struc** | **NetSurfP predicted probability score of the three secondary structure types** |
| PA_Q | score for $\alpha$ helix for the query sequence |
| mean_PA, PA_std | average and standard deviation score for $\alpha$ helix for each column in the alignment |
| PB_Q | score for $\beta$ sheet for the query sequence |
| mean_PB, PB_std | average and standard deviation score for $\beta$ sheet for each column in the alignment |
| PC_Q | score for coil for the query sequence |
| mean_PC, PC_std | average and standard deviation score for coil for each column in the alignment |
| **Conservation** | **Sequence entropy** |
| H_Entropies | Sequence entropy used to describe the degree of conservation of the query sequence, based on the PSI-BLAST profile |

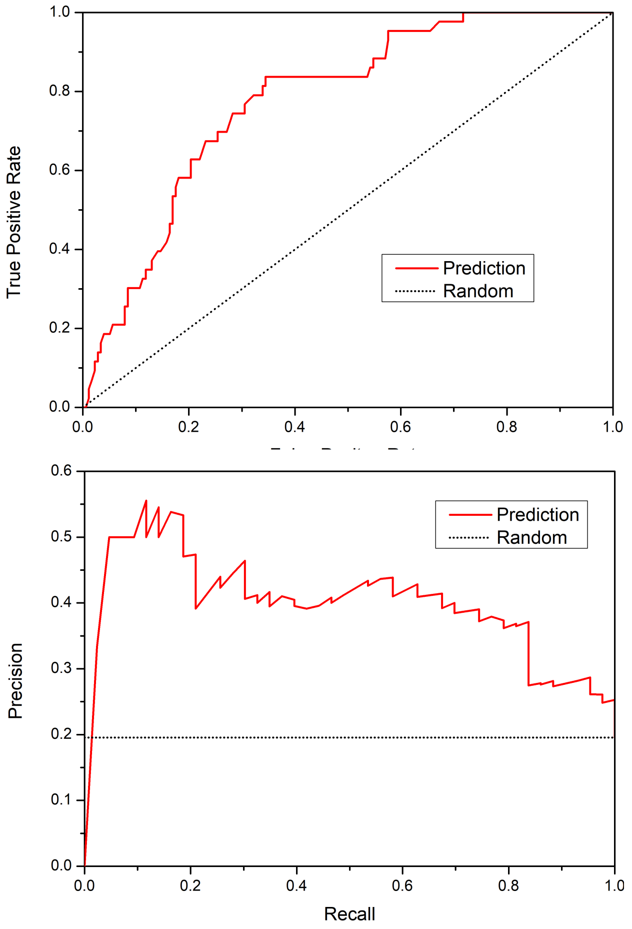


**SI Figure S4.** The average performance of our predictors on COVID19 RBD domain 7BZ5:A. The ROC plot and the Precision-Recall plot are shown here

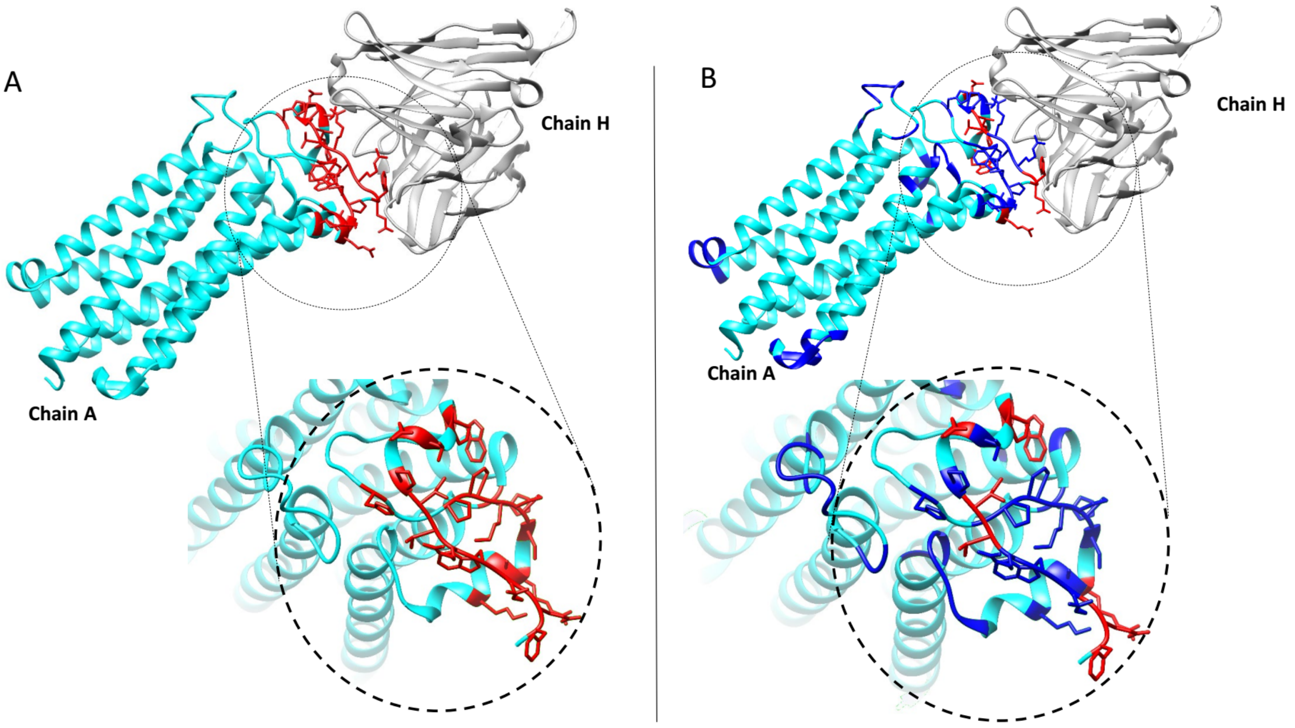


**SI Figure S5.** Epitope prediction on Adiponectin receptors-antibody complex We show one example of epitope predictions using Adiponectin receptors-2 as antigen.The antigen is shown in cyan, epitopes in red. (A) the whole structure of antibody-antigen and the zooming details of the epitope interface. (B) The prediction of our approach using default cut-off 0.5. The positions with the probability score higher than 0.5 are shown in blue.

To highlight the impact of accurate epitope prediction, we here show an example. Adiponectin receptors (ADIPORs) are integral membrane proteins that control glucose and lipid metabolism. The crystallization of Adiponectin receptors-2 (PDB ID 5LX9) was performed with antibody fragments and a protein fusion, in the lipidic mesophase. We applied the predictor on the Adiponectin receptor sequence as antigen input. The protein was not part of our training set, and its sequence identity to any protein in our training data is less than 25%. The predictions and real interface are shown in the crystal structure of the complex. For this particular interaction, a coverage of 58% of the 24 interface sites at an accuracy of 82% is achieved, yielding an AUC-ROC of 0.819. As can be seen from panel B, our predictions cover the majority of the epitope positions. Interestingly, our false positive predictions clustered on the other side of protein which might be due to the fact that the other side of trans membrane protein is also amenable to antibody binding.
